# Comprehensive simulation and interpretation of single nucleotide substitutions in *GJB2* reveals the genetic and phenotypic landscape of *GJB2*-related hearing loss

**DOI:** 10.1101/2021.09.30.462500

**Authors:** Jiale Xiang, Xiangzhong Sun, Nana Song, Lisha Chen, Sathishkumar Ramaswamy, Ahmad Abou Tayoun, Zhiyu Peng

## Abstract

Genetic variants in the *GJB2* gene are the most frequent causes of congenital and childhood hearing loss worldwide. In addition to nonsyndromic hearing loss, *GJB2* pathogenic variants are also correlated with syndromic phenotypes, showing high genetic and phenotypic heterogeneity. To comprehensively delineate the genetic and phenotypic landscape of *GJB2* variants, we interpreted and manually curated all the 2043 possible single-nucleotide substitution (SNS) coding variants in this gene following the hearing loss-specific ACMG/AMP guidelines. As a result, 61 (3.0%), 188 (9.2%), 1487 (72.8%), 301 (14.7%) and 6 (0.3%) variants were classified as pathogenic, likely pathogenic, variant of uncertain significance, likely benign and benign, respectively. Interestingly, 54% (84/156) of pathogenic/likely pathogenic missense variants were not recorded in ClinVar. Further analysis showed that the second transmembrane domain (TM2) and the 3_10_ helix are highly enriched for pathogenic missense variants. The N-terminal tail and the extracellular loop (E1) showed a high density of variants that are associated with syndromic or dominant nonsyndromic hearing loss. On the other hand, the intracellular loops (CL and CT) were extremely tolerant to variation. Based on this new information, we propose refinements of the guidelines for variant interpretation in *GJB2*. In summary, our study interpreted all possible SNS variants in the coding region of the *GJB2* gene, characterized novel clinically significant (N = 249) and benign or likely benign (N = 307) in this gene, and revealed significant genotype-phenotype correlations at this common hearing loss locus. The interpretation of *GJB2* SNS variants in the coding region provides a prototype for genes with similarly high genetic and phenotypic heterogeneity.

## Introduction

The prevalence of congenital hearing loss (HL) is 1-3 infants per 1,000 live births, making it one of the most common congenital disorders worldwide.^1^ Although the causes are multifactorial, over half of which is genetic in etiology.^1^ Hearing loss presents extraordinary genetic and phenotypic heterogeneity. To date, more than 400 syndromes associated with hearing loss have been described, and more than 140 genes associated with nonsyndromic hearing loss have been identified.^2; 3^

Gap junction protein β2 (*GJB2*), the most prevalent deafness-related gene, is responsible for nearly 50% of profound nonsyndromic hearing loss cases in many populations.^4^ The *GJB2* gene, located on chromosome 13q12.11, has a 2290-nucleotide mRNA (GeneBank: NM_004004.6), which is composed of two exons, only one of which (exon 2) is coding (681-nucleotide coding region). Connexin 26 (Cx26), encoded by the *GJB2* gene, is widely expressed in supporting cells and connective tissues of the cochlea, playing a role in potassium ion recycling and inositol triphosphate transfer.^5; 6^ Cx26 consists of four α-helical transmembrane domains (TM1–TM4), two extracellular loops (E1 and E2), a cytoplasmic loop (CL) between TM2 and TM3, and cytoplasmic amino-terminal (NT) and carboxy-terminal (CT) domains.^7^ Six connexin protomers oligomerize to form a transmembrane connexon, and two connexons in adjacent cells align end-to-end to form a complete intercellular gap junction channel.^7^

Pathogenic variants in the *GJB2* gene cause a variety of phenotypes. Autosomal recessive nonsyndromic hearing loss (DFNB1A, OMIM #220290) is the most common one. It is contributed by the loss-of-function of mutant Cx26, impairing protein folding, trafficking, oligomerization or gap junction channel formation.^8^ On the other hand, gain-of-function variants generally lead to formation of leaky hemichannels, hyperactive connexons and/or gap junctions.^9^, and result in autosomal dominant disorders with or without syndromic skin disorders, including autosomal dominant nonsyndromic hearing loss (DFNA3A, OMIM #601544), keratitis-ichthyosis-deafness syndrome (KID syndrome, OMIM #148210), palmoplantar keratoma with deafness (PPK syndrome, OMIM #148350), hystrix-like ichthyosis with deafness (HID syndrome, OMIM #602540), Bart-Pumphrey syndrome (OMIM #149200) and Vohwinkel syndrome (OMIM #124500). Considering its major contribution to hearing loss along with the associated genetic and phenotypic heterogeneity, it is of significant clinical and research priority to delineate the comprehensive genetic and phenotypic landscape of *GJB2* in hearing loss.

To fulfill the goal, we manually curated all possible single-nucleotide substitution (SNS) variants in the coding region of *GJB2*, following the clinically adopted variant interpretation guidelines for genetic hearing loss.^10^ We analyzed the distribution of missense variants by pathogenicity and genotype-phenotype correlations across the protein structure of Cx26. Based on our data, we refined the variant interpretation criteria for this gene, and estimated the prevalence of DFNB1A in different ethnicities.

## Material and Methods

### SNS Variants and Variant Annotation

The *GJB2* gene has a 681-nucleotide coding region. Given three substitutions for each nucleotide, there are 2043 single-nucleotide substitution variations in the coding region of *GJB2* (Figure 1A). All SNS variants were described at DNA level and protein level according to Human Genome Variation Society nomenclature.^11^ Additionally, variants were annotated with consequences (including missense, synonymous, nonsense, start loss, or stop loss) via Ensembl Variant Effect Predictor.^12^ The highest filtering allele frequency (AF) across all populations (popmax filtering allele frequency) were acquired from Genome Aggregation Database (gnomAD, v2.1).^13; 14^ ClinVar classifications for each variant were retrieved from ClinVar 20210204 release.^15^ Variants that are included in Human Gene Mutation Database (HGMD, 2019.2 release)^16^ and Deafness Variation Database (DVD, v9)^17^ were also annotated.

**Figure 1.**
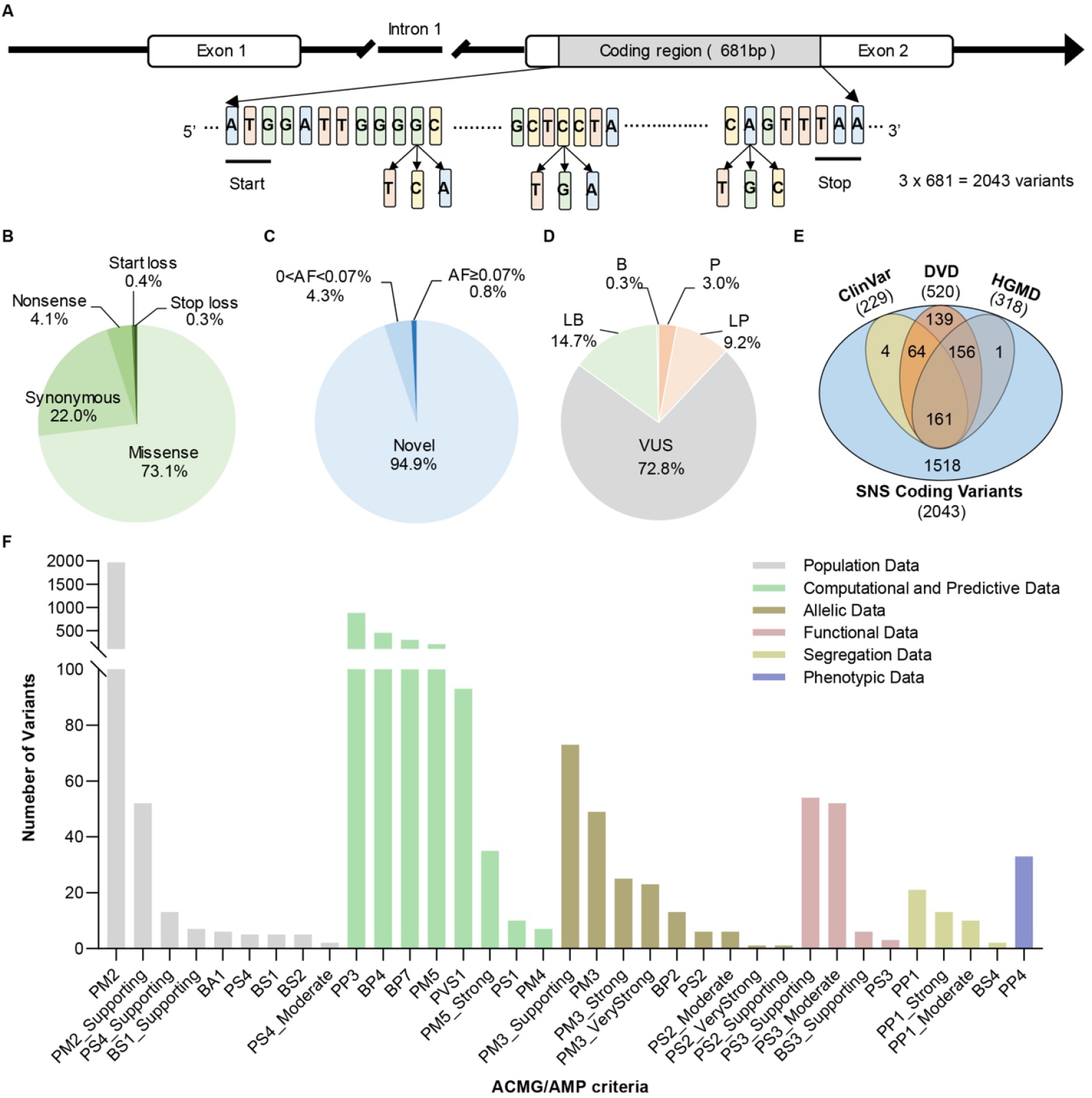
The characteristics of the *GJB2* gene and 2043 single-nucleotide substitution variants on the coding region. (A) A schematic presentation of the *GJB2* gene structure and SNS variants on the coding region. (B) The predicted consequence of 2043 SNS variants. (C) The allele frequency of 2043 SNS variants in the gnomAD database. (D) The classifications of 2043 SNS variants based on the expert specification of the ACMG/AMP variant interpretation guidelines for genetic hearing loss. (E) Venn diagram of SNS variants in ClinVar, HGMD, DVD and this study. (F) Activated ACMG/AMP criteria for 2043 SNS variants by type. Abbreviation: P, pathogenic; LP, likely pathogenic; VUS, Uncertain Significance; LB, likely benign; B, benign; SNS, single-nucleotide substitution; gnomAD, Genome Aggregation Database; HGMD, Human Gene Mutation Database; DVD, Deafness Variation Database; ACMG/AMP, the American College of Medical Genetics and Genomics/Association for Molecular Pathology.

### Variant Interpretation and Manual Curation

All the variants were curated based on the expert specification of the American College of Medical Genetics and Genomics/Association for Molecular Pathology (ACMG/AMP) variant interpretation guidelines for genetic hearing loss.^10^ Of 24 ACMG/AMP criteria, population (PM2, BA1, BS1 and BS2) and computational (PVS1, PM4, PP3, BP3, BP4, and BP7) data criteria were automatically curated by Variant Interpretation Platform for Genetic Hearing Loss (VIP-HL).^18; 19^ Criteria related to allelic data (PM3, PS2/PM6 and BP2), functional data (PS3 and BS3), segregation data (PP1 and BS4), phenotypic (PP4 and BP5) and case/control (PS4) data were manually curated from public literature by experienced biocurators. Considering that there are no mutational hot spots or well-studied functional domains without benign variation in *GJB2*,^10^ PM1 was not activated. The final pathogenicity is reported in five tiers system, namely pathogenic (P), likely pathogenic (LP), variant of uncertain significance (VUS), likely benign (LB), and benign (B).^20^ Interpreting discrepancies were discussed biweekly. If no agreement was reached, the variants were sent for senior researchers (Dr. Peng and Dr. Abou Tayoun) for further discussion. The final classifications were reviewed by a multidisciplinary expert panel including clinicians, geneticists, bioinformaticians and genetic counselors.

### Comparative Analysis

ClinVar was chosen for comparative analysis. In detail, all SNS variants on the coding region of *GJB2* were retrieved from ClinVar 20210204 release. The ClinVar star 2+ (i.e., multiple submitters with assertion criteria, expert panel, or practice guideline) variants were separately compared because these variants had fewer misclassifications.^21^ The comparison was analyzed in three tiers amongst P/LP, VUS and B/LB. If the differences did not affect clinical interventions, they were regarded as medical concordance (i.e., P vs. LP or B vs. LB). Because P/LP classifications are clinically actionable, we calculated medically significant differences when variants were downgraded from or upgraded to P/LP (i.e., P/LP vs. VUS).

### Estimated Population-specific Prevalence of DFNB1A

The estimated prevalence of DFNB1A was calculated in eight ethnicities. The allele frequency was retrieved from gnomAD^13^, except for the Middle East, whose genome was less representative in gnomAD.^22^ The allele frequency of the Middle East was retrieved from our in-house data.^23^ Assuming that P/LP variants are in linkage equilibrium and the eight populations are in Hardy-Weinberg equilibrium (HWE), we estimated the cumulative genetic risk (GR) for homozygous and compound heterozygous pathogenic recessive variants (DFNB1A) as follows:^24^

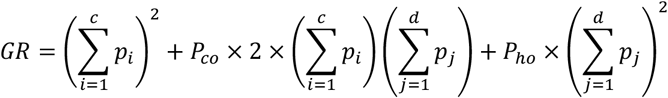

where *p* is the allele frequency for each variant, *c* is the number of P/LP variants with complete penetrance, *d* is the number of P/LP variants with incomplete penetrance, *P*_*ho*_ is the penetrance of homozygotes with the partial penetrant P/LP variants, and *P*_*co*_ is the penetrance of corresponding compound heterozygotes.

For each population, the recessive P/LP variants are sorted and arranged in the order of allele frequency. Then, the cumulative genetic risk was calculated with the different number of variants sequentially, starting from the highest allele frequency variant within the population.

In this study, all the *GJB2* recessive P/LP variants were assumed with complete penetrance, except for c.101T>C (p.Met34Thr) and c.109G>A (p.Val37Ile), which have been reported to be associated with incomplete penetrance.^25; 26^ The penetrance of compound heterozygotes of these two variants *in trans* with other *GJB2* pathogenic variants is set as 10%,^25^ and the penetrance of homozygotes for either variant is set as 5%, lower than that in the corresponding compound heterozygotes.^26^

## Results

### Overview of the 2043 *GJB2* SNS Coding Variants

Considering three single substitutions for each nucleotide, a total of 2043 SNS variants were calculated from the 681 bp coding region of the *GJB2* gene (Figure 1A). The majority of the SNS variants were missense (1493, 73.1%), followed by synonymous (450, 22.0%), nonsense (84, 4.1%) and start/stop loss (16, 0.8%) (Figure 1B and Table S1). Of all SNS variants, novel (AF = 0) and ultra-rare (0% < AF < 0.07%) variants represented 94.9% and 4.3%, respectively. Variants with an AF ≥ 0.07% (the threshold which benign criteria can be applied to a variant in genetic hearing loss^10^) represented only 0.8% (Figure 1C and Table S1).

Applying the expert specification of the ACMG/AMP variant interpretation guidelines for genetic hearing loss, 61 (3.0%), 188 (9.2%), 1487 (72.8%), 301 (14.7%) and 6 (0.3%) variants were classified as P, LP, VUS, LB and B, respectively (Figure 1D). The ACMG/AMP criteria for all 2043 variants were listed in Table S1. Of 2043 SNS variants, 1518 (73.3%) were not recorded in ClinVar, DVD, and HGMD (Figure 1E). More importantly, the majority (61%, 151/249) of P/LP variants were not recorded in ClinVar (Figure S1). The 249 P/LP variants comprised 156 missense variants and 93 nonsense or start loss variants. 54% (84/156) and 72% (67/93) of P/LP missense and nonsense or start loss variants, respectively, were not recorded in ClinVar (Figure S1). For the 84 non-ClinVar P/LP missense variants, criteria related to functional data, allelic data, and phenotypic data (PS3, PM3, PP4, etc.) were intensively activated (Figure S2), indicating that these variants are clinically reportable and medically relevant.

The frequencies of activated ACMG/AMP criteria were analyzed (Figure 1F and Table S1). A total of 4434 criteria were activated for 2043 variants, with an average of 2.17 criteria per variant. The population frequency codes were most frequently applied, following by computational and predictive data (Figure 1F). The allelic, functional, segregation, and phenotypic data were manually curated from the literature and were relatively less frequently activated. Of note, PM3 and its modified strength levels were activated for 170 (8.3%) variants; functional evidence (PS3, BS3 and their modified strength levels) were activated for 109 (5.5%) variants. PP1 and its modified strength levels (segregation data) and PP4 (phenotypic data) were only activated for 44 (2.2%) and 33 (1.4%) variants, respectively. These results exhibited the diverse and extensive activations of ACMG/AMP criteria in the interpretation of variants in the *GJB2* gene.

### Comparison with ClinVar Classification

To assess the interpreting consistency between ClinVar and this study, we compared medical concordance and medically significant differences (upgraded to or downgraded from P/LP) in the two resources. Of the 2043 SNS variants, 229 were found in ClinVar (Figure 1E). By comparison, 187 (82%) variants had medical concordance, whereas 27 (12%) had medically significant differences (Figure 2). Specifically, 17 variants were downgraded from P/LP to VUS; 10 variants were upgraded from VUS to P/LP. The comments for upgrade and downgrade were listed in Table S2.

**Figure 2.**
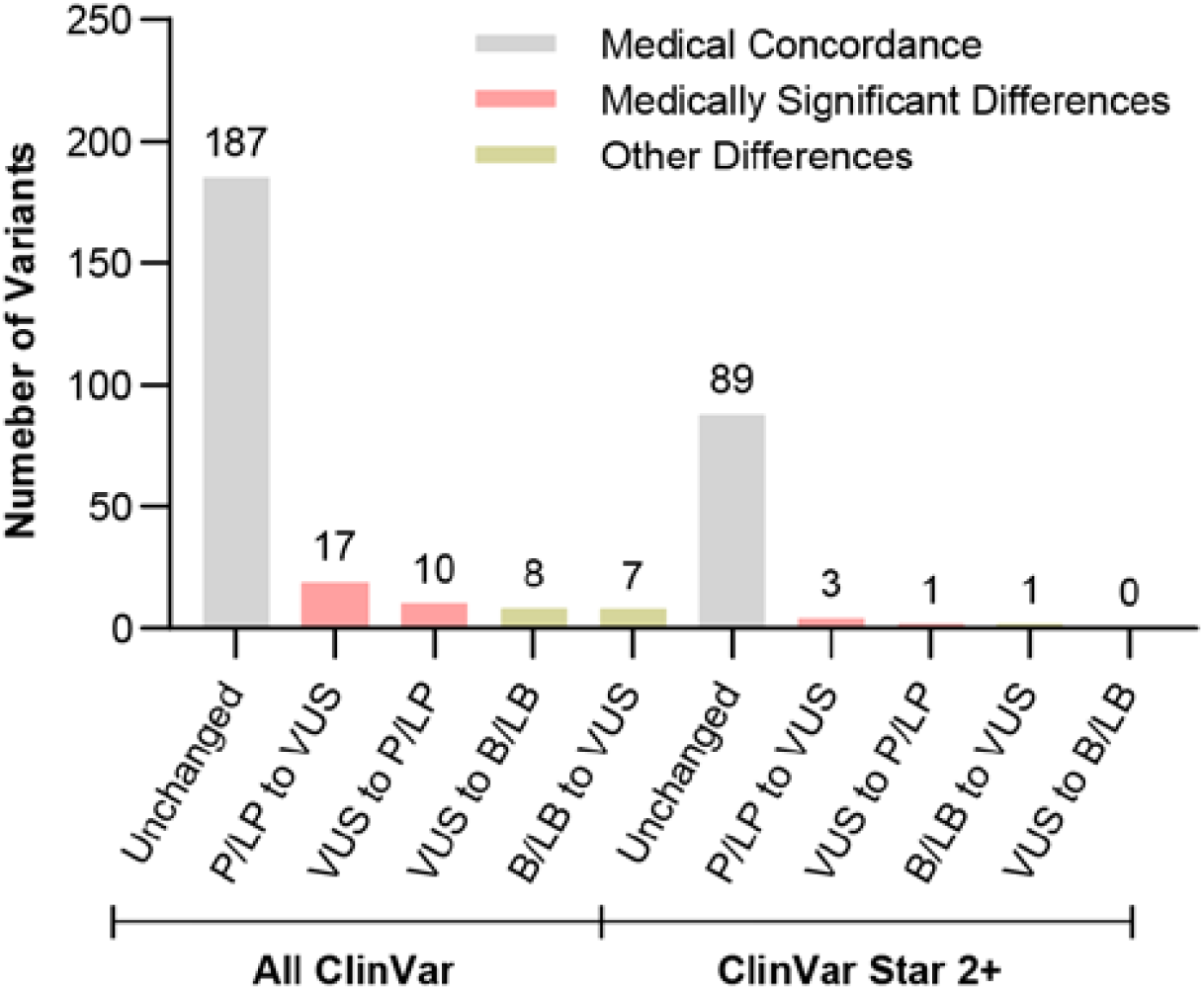
Distribution of variant interpretation differences between ClinVar and this study. Abbreviation: P, pathogenic; LP, likely pathogenic; VUS, variant of uncertain significance; LB, likely benign; B, benign.

Of all these 229 SNS variants in ClinVar, 94 variants are assigned two or more gold stars. Overall, 89 (95%) 2-star variants had medical concordance, 4 (4%) had medically significant differences, 1 (1%) had other differences (Figure 2). Of the five discordant variants, c.34G>T (p.Gly12Cys) and c.110T>C (p.Val37Ala) have been reviewed by ClinGen Hearing Loss Expert Panel (HL-EP). c.34G>T (p.Gly12Cys) was classified as LP based on PM3_VeryStrong, PM5, PP3, BS1; c.110T>C (p.Val37Ala) was classified as LP based on PM3, PM5, PM2_Supporting, PP3. Notably, internal allelic data from Laboratory for Molecular Medicine (Partners HealthCare) were used for the interpretation of both variants, which were not available for this study. Without the activations of PM3, the two variants would be classified as VUS, which was consistent with our classification. The remaining three variants were submitted to ClinVar by multiple submitters without the provision of ACMG/AMP criteria. Based on our interpretation, c.533T>C (p.V178A) was downgraded from P/LP to VUS because the available information was not sufficient to reach a P/LP classification. c.584T>C (p.M195T) was upgraded from VUS to LP based on PS3_Moderate, PM2, PM5, PP3. The last variant, c.412A>G (p.S138G), was upgraded from B/LB to VUS due to a lack of benign evidence (Table S2).

Overall, the comparison and analysis demonstrated that the classifications of 2043 variants in this study, and our approach, are reliable.

### The Landscape of *GJB2* Missense Variants

Having built a comprehensive list of all 2043 SNS coding variants in the *GJB2* gene with clinical interpretations, we sought to explore the intragenic localization and any genotype-phenotype correlations of *GJB2* missense variants. We analyzed the distribution of the pathogenicity in nine structural motifs of Cx26 (Figure 3 and Table 1).

**Figure 3.**
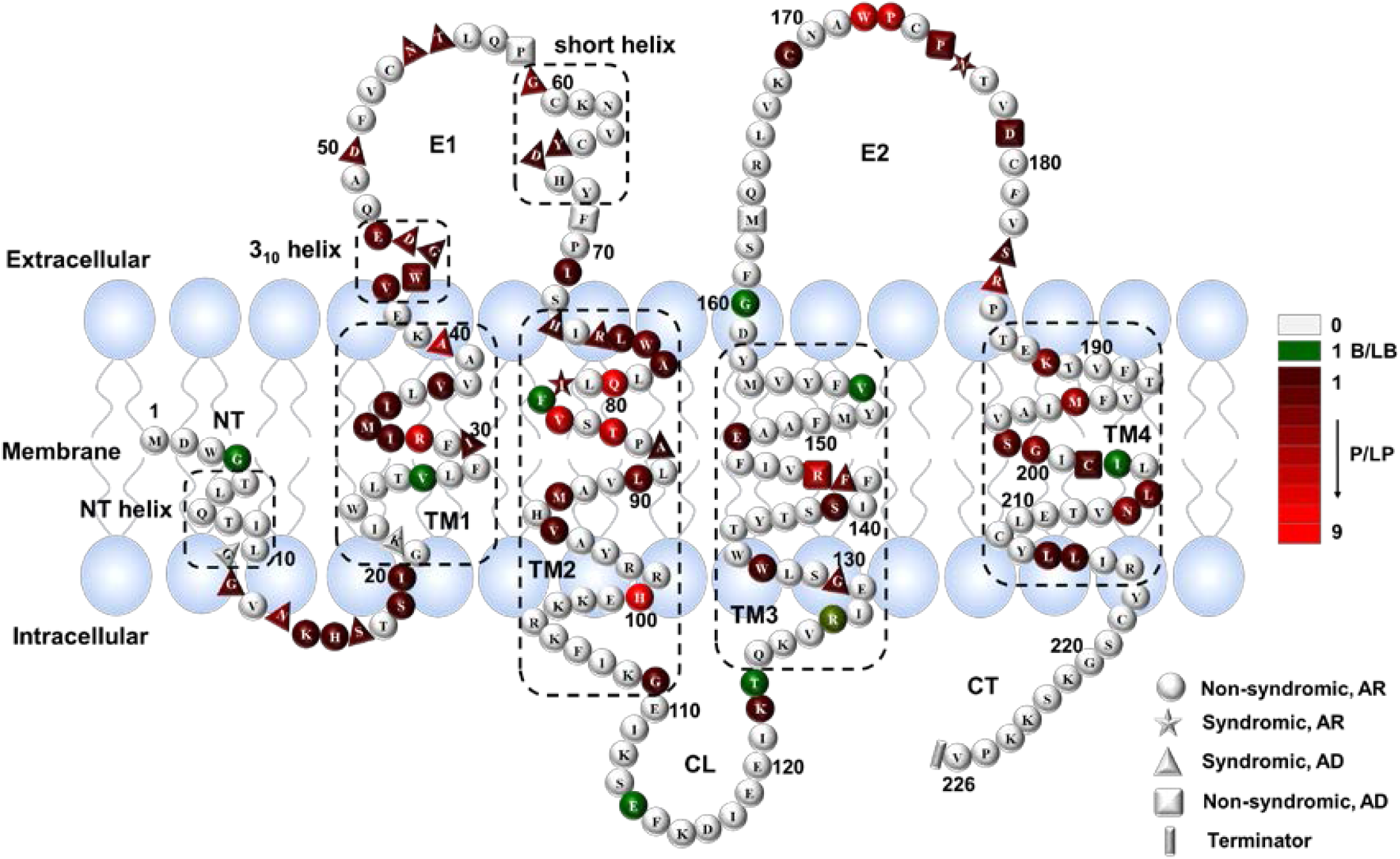
Topological diagram of the Cx26 protomer. The P/LP or B/LB missense variants are indicated by color. The genotype and inheritance are indicated by shapes or symbols. Dotted boxes indicate regions with a helical secondary structure denoted as NT helix, TM1, TM2, TM3, TM4, 3_10_ helix and short helix in E1. Abbreviation: P, pathogenic; LP, likely pathogenic; LB, likely benign; B, benign; AR, autosomal recessive; AD, autosomal dominant. TM1-TM4, four transmembrane domains; E1 and E2, two extracellular loops; NT, N-terminal tail; CL, cytoplasmic loop; CT, carboxy-tail.

**Table 1.** The distribution of pathogenicity of *GJB2* missense variants on nine structural motifs.

| Structural Motif | aa Length (aa range) | Number of Missense Variants |  |  |  | Pathogenic Rate (%)<sup>a</sup><br>(AR variants only) | Normalized No. of P/LP<sup>b</sup><br>(AR variants only) |
| --- | --- | --- | --- | --- | --- | --- | --- |
|  |  | B/LB | VUS | P/LP<br>(AR variants only) | Total |  |  |
| NT | 20 (1-20) | 1 | 113 | 11 (6) | **125** | 9% (5%) | 0.55 (0.30) |
| TM1 | 21 (21-41) | 1 | 121 | 17 (15) | **139** | 12% (11%) | 0.81 (0.71) |
| E1 | 31 (42-72) | 0 | 186 | 24 (7) | **210** | 11% (3%) | 0.77 (0.22) |
| TM2 | 37 (73-109) | 1 | 196 | 43 (39) | **240** | 18% (16%) | 1.16 (1.05) |
| CL | 14 (110-123) | 2 | 97 | 1 (1) | **100** | 1% (1%) | 0.07 (0.07) |
| TM3 | 35 (124-158) | 2 | 221 | 14 (10) | **237** | 6% (4%) | 0.40 (0.28) |
| E2 | 27 (159-185) | 1 | 154 | 25 (20) | **180** | 14% (11%) | 0.93 (0.74) |
| TM4 | 31 (186-216) | 1 | 178 | 21 (20) | **200** | 11% (10%) | 0.68 (0.65) |
| CT | 10 (217-226) | 0 | 62 | 0 (0) | **62** | 0% (0%) | 0.00 (0.00) |
| **Total** | **226** | **9** | **1328** | **156 (118)** | **1493** | **10% (8%)** | **0.69 (0.52)** |

**a**, The rate of pathogenic/likely pathogenic missense variants on each motif. (Pathogenic Rate = $\frac{\text{No.of P/LP missense variants}}{\text{No.of all missense variants}}$ )
**b**, Normalized number of pathogenic/likely pathogenic missense variants based on the length of motif. (Normalized No. of P/LP = $\frac{\text{No.of P/LP missense variants}}{\text{amino acid length}}$ )
Abbreviation: **TM1-TM4**, four transmembrane domains; **E1** and **E2**, two extracellular loops; **NT**, N-terminal tail; **CL**, cytoplasmic loop; **CT**, carboxy-tail. aa, amino acid.

Clearly, the P/LP variants were greatly enriched in TM2; the pathogenic rate was 18% (43/240), 1.8 times higher than the average rate (10%, 156/1493; P<0.001; hypergeometric test). If only the autosomal recessive variants were calculated, the pathogenic rate was 16% (39/240), indicating most P/LP variants in TM2 were inherited recessively (Table 1). Moreover, P/LP variants in TM2 were not equally distributed; most of those variants were clustered in the TM2 region proximal to the E1 domain (Figure 3).

Interestingly, the 3_10_ helix, a five-residue segment spanning V43–E47 in the Cx26 crystal structure,^7^ was also enriched with pathogenic missense variants (Figure 3). More specifically, the autosomal dominant variants were highly enriched (P = 0.00014, hypergeometric test; Table S3). The results strongly suggested that the 3_10_ helix is critical for Cx26 function. From the perspective of variant interpretation, considering the enrichment of P/LP variants and absence of benign missense variants, we recommend activating PM1 for a novel rare missense variant occurring in the 3_10_ helix segment in association with dominant phenotype. By applying this rule, the ratio of VUS missense variants on the 3_10_ helix was decreased from 70% (23/33) to 58% (19/33).

In addition, CT and CL are the most tolerant to variation. More specifically, no P/LP variant was reported on the CT motif. Only 1 (1%) LP variant was reported on the CL motif. The evidence prompts us to consider CL and CT of Cx26 as protein “Cold spots”, describing regions of a gene that are tolerant to variation, where pathogenic missense variants are unlikely.^27^ Thus, missense variants on CL and CT could be counted as benign piece of evidence.

It was obvious that the pathogenic rate of missense variants on NT and E1 differed when only the autosomal recessive variants were calculated (Table 1), indicating that a high proportion of missense variants on these two motifs are inherited dominantly. To examine this hypothesis, we listed all dominant missense variants by phenotypes and structural motifs (Table 2). Of 40 missense variants inherited dominantly, 17 (43%) were located on E1. Variants located at the E1/TM1 boundary (A40V) or E1/TM2 boundary (H73R, R75W, R75Q) were also reported to exhibit syndromic deafness. Besides, variants affecting four residues (G11, G12, N14 and S17) on the NT motif are all associated with autosomal dominant KID syndrome. To date, none of the variants with syndromic and dominant nonsyndromic phenotypes have been reported on the cytoplasmic loops (CL and CT) (Figure 3 and Table 2). It is worth noting that although syndromic phenotypes are generally inherited in an autosomal dominant manner in the *GJB2* gene, two variants (I82V and N176D) causing syndromic phenotypes were reported to segregate recessively.^28; 29^ These results shed light on the phenotypic heterogenicity of Cx26, and the distribution of disease-causing missense variants within *GJB2*.

**Table 2.** The genotype-phenotype (syndromic and dominant non-syndromic hearing loss) associations of missense variants in the *GJB2* gene.

| Phenotype | OMIM number | Structural Motif |  |  |  |  |  |  |  |  |
| --- | --- | --- | --- | --- | --- | --- | --- | --- | --- | --- |
|  |  | NT | TM1 | E1 | TM2 | CL | TM3 | E2 | TM4 | CT |
| **DFNA3A** | 601544 | - | - | W44C, D46N, **D46E**, P58A, F69L | - | - | G130V | M163V, P175H, D179N | C202F | - |
| **KID syndrome** | 148210 | G11E, G12R, N14Y, N14K, S17F | A40V | G45E, **D50N**, D50Y, D50A | I82V<sup>#</sup>, A88V | - | F142L | P175R | - | - |
| **PPK syndrome** | 148350 | - | K22N, I30N | **D46E**, N54H, T55A, G59A, **G59R** | H73R, R75W, R75Q | - | - | N176D<sup>#</sup>, S183F, R184Q | - | - |
| **Vohwinkel syndrome** | 124500 | - | - | **G59R**, G59S, Y65H, D66H | - | - | - | - | - | - |
| **Bart-Pumphrey syndrome** | 149200 | - | - | N54K | - | - | - | - | - | - |
| **HID syndrome** | 602540 | - | - | **D50N** | - | - | - | - | - | - |

### Refinement of Variant Interpretation in *GJB2*

The *GJB2* gene exhibited extraordinary genetic and phenotypic heterogeneity, which posed great challenges for variant interpretations and curations. Of 236 missense variants with established inheritance patterns, 191 (81%) were autosomal recessive and 43 (18%) were autosomal dominant (Table S4). In this study, we encountered two new scenarios relevant to variant interpretation in *GJB2*. The first scenario was a variant inherited both recessively and dominantly. Two missense variants, c.205T>C (p.F69L) and c.428G>A (p.R143Q), were reported to exhibit both recessive and dominant inheritance patterns (Table S4 and S5). The two variants were curated for different inheritance patterns separately; that is, ACMG/AMP criteria supporting autosomal recessive and dominant manner did not apply simultaneously for curations. For example, c.205T>C (p.F69L) was reported *in trans* with c.35delG in a proband with hearing loss (PM3),^30^ implying an autosomal recessive manner. Huang et al. reported that this variant co-segregated with prelingual nonsyndromic sensorineural hearing loss in a dominant pattern, supporting the activation of PP1_Moderate.^31^ As a result, we applied PM3, PM2, PP3 for c.205T>C for autosomal recessive, and PP1_Moderate, PM2, PP3 for autosomal dominant. The final classifications for both inheritance patterns were VUS, which is different from the pathogenic classification in ClinVar (Table S5).

The second scenario is accounting for both inheritance patterns on the same amino acid residue. Of all the 158 amino acid residues with established inheritance patterns, autosomal recessive, autosomal dominant, and dual mode of inheritance presented on 128 (81%), 9 (6%), and 21 (13%) amino acid residues, respectively (Table S4). The amino acid change with different inheritance patterns leads to a distinct phenotype. This phenotypic heterogenicity with variable inheritance patterns on the same amino acid affects the activation of PM5, a scenario which has not been explicitly addressed by previous guidelines^10; 20^. Therefore, we interrogated all the 1493 missense variants in the *GJB2* gene with respect to inheritance pattern and PM5 activation (Table S6). Interestingly, the LP variants were mainly impacted by this notion. With and without the consideration of inheritance patterns, the number of LP variants was 124 and 162, respectively. That is, 38 variants (31%, 38/124) had medically significant differences (VUS vs. LP) when the inheritance pattern was taken into consideration while activating PM5. Based on our data, we proposed a decision tree to specify the application PM5 (Figure 4). PM5 can be activated when the inheritance pattern of a novel variant (variant being interpreted) and the established pathogenic variant is consistent. Otherwise, PM5 should not be applied.

**Figure 4.**
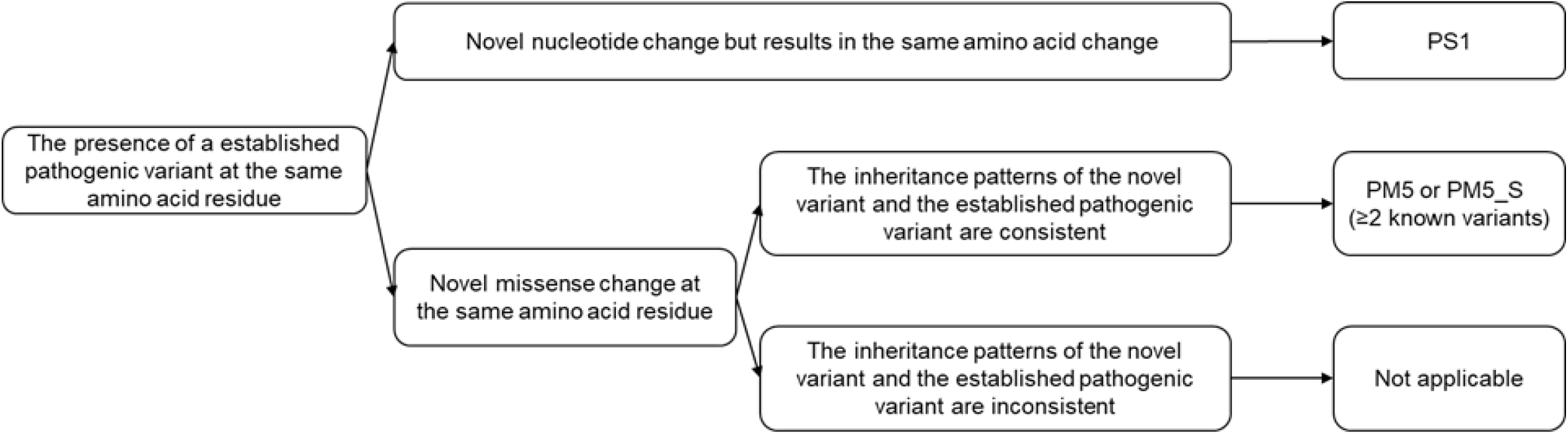
Recommended PS1/PM5 decision tree.

### Expected Prevalence of DFNB1A Caused by *GJB2* Pathogenic Variants

*GJB2*-related mutations are the most common etiology in congenital nonsyndromic hearing loss. With the availability of high reliable classifications for variants in the *GJB2* gene, we sought to explore the estimated prevalence of DFNB1A in different populations. Considering the prevalence of congenital hearing loss is 0.1% ∼ 0.3% in clinical settings, and about 20% of congenital hearing loss is caused by *GJB2* mutations,^32^ the observed prevalence of DFNB1A is calculated to 2 × 10^−4^ ∼ 6 × 10^−4^ (Figure 5). The estimated prevalence was calculated through Hardy-Weinberg equilibrium (See Methods). Of 252 recessive P/LP variants, 83 variants were observed in at least one population in gnomAD and 169 were novel (Table S7 and S8). The allele frequencies of recessive P/LP variants displayed considerable variability between populations, leading to the highly variable estimated prevalence of DFNB1A amongst eight populations (Figure 5 and S7). East Asian population had the highest estimated prevalence (5.72 × 10^−4^) and African American population had the lowest (1.76 × 10^−5^). The estimated prevalence for other populations was 4.97 × 10^−4^ (Ashkenazi Jewish), 2.31 × 10^−4^ (Non-Finnish European), 1.43 × 10^−4^ (Finnish), 1.11 × 10^−4^ (Latino), 7.11 × 10^−5^ (South Asian), and 2.11 × 10^−5^ (Middle East), respectively.

**Figure 5.**
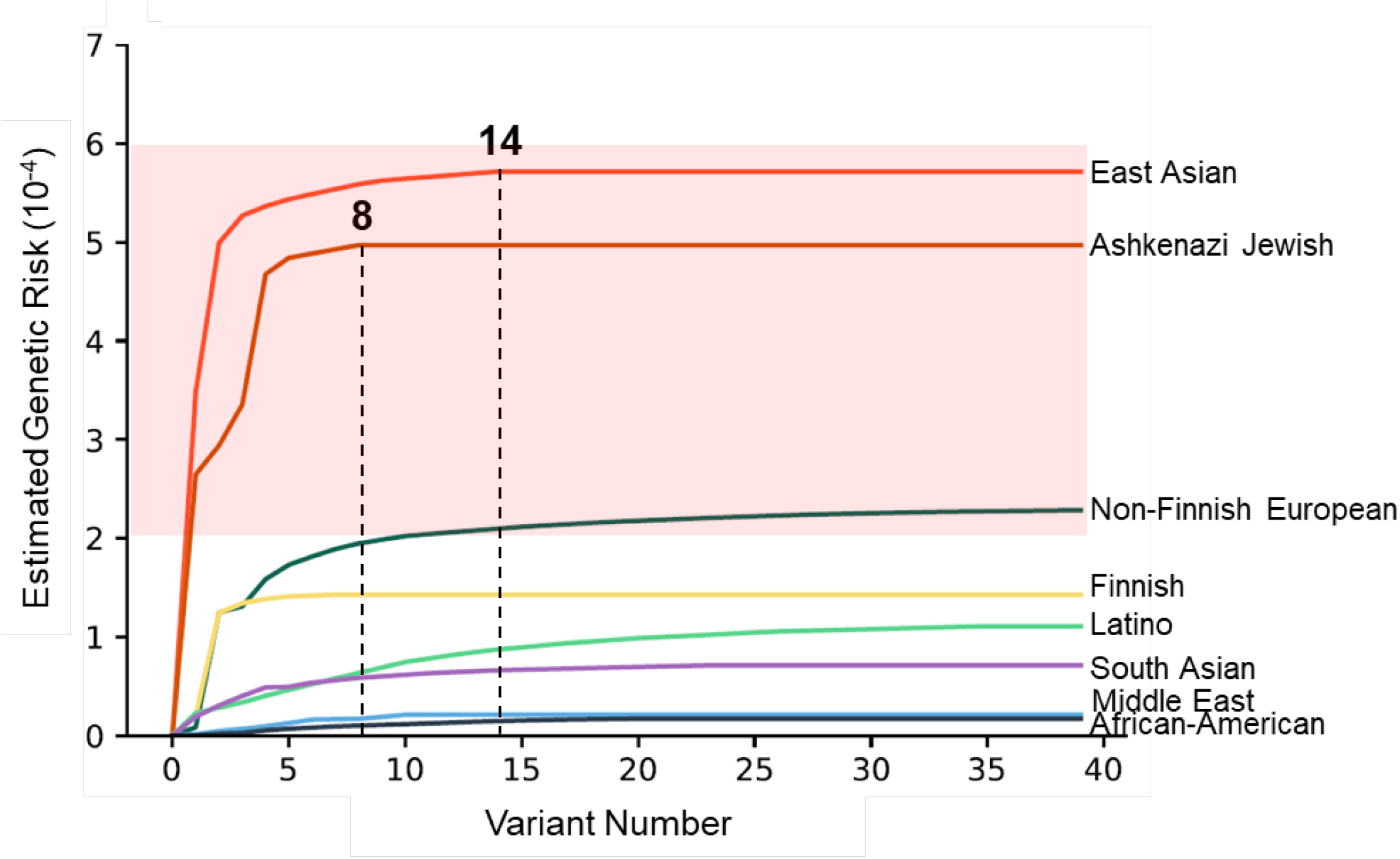
The cumulative genetic risk of DFNB1A in eight ethnicities. The cumulative genetic risk of DFNB1A was calculated based on Hardy-Weinberg equation. The observed prevalence of DFNB1A is indicated in a shade of pink. For East Asian and Ashkenazi Jewish, the total numbers of recessive pathogenic/likely pathogenic variants observed in gnomAD are indicated.

Next, we analyzed the contribution of variants to the estimated prevalence. For East Asian, only 14 P/LP variants had allele frequency in gnomAD, and 95% of the estimated prevalence was driven by the top five variants (c.109G>A, c.235del, c.299_300del, c.508_511dup, c.583A>G). Ashkenazi Jewish population had the second-highest genetic risk of DFNB1A (4.97 × 10^−4^), with only 8 P/LP variants were observed in gnomAD. The top five variants (c.167del, c.109G>A, c.101T>C, c.35del, c.416G>A) accounted for 97% estimated prevalence of DFNB1A. Non-Finnish Europeans had the highest diversity of recessive P/LP variants, counting 48 variants in gnomAD. Despite the high diversity, the first 13 variants accounted for 90% estimated prevalence. These findings demonstrated that *GJB2* pathogenic variants causing DFNB1A is highly variable between populations. More importantly, the high prevalence was contributed by a limited number of variants. This analysis can be the foundation to guide the much-needed pan-ethnic newborn screening for *GJB2* variants.

## Discussion

Genetic variant interpretation plays a significant role in accurate genetic diagnosis, particularly for a disease with high genetic and phenotypic heterogeneity like hearing loss. In this study, we interrogated massive information from public databases, literature, bioinformatics tools, then comprehensively interpreted all 2043 SNS variants on the coding region of the *GJB2* gene, the most common cause of genetic hearing loss. All the classifications are based on clinically adopted variant interpretation guidelines for genetic hearing loss.^10^ To our knowledge, this is the first time that the genetic and phenotypic landscape of Cx26 is illustrated. It is also the first gene to be comprehensively analyzed using our approach. Considering the high prevalence of congenital and childhood hearing loss and the contribution of *GJB2* to the etiology, our work provides a significant resource for clinicians, genetic professionals, genetic counselors, and patients.

Of 2043 SNS variants, 12.2% (249/2043) were classified as P/LP, indicating every one in eight SNS variants on the coding region of the *GJB2* gene is clinically significant. In comparison, ∼46% of SNS coding variants of the *GJB2* gene are recorded as P/LP in ClinVar (Figure S3); ∼69% of coding variants in *GJB2* are P/LP in DVD.^17^ The ratio is lower in this study because all the variants were manually and conservatively curated based on the expert specification of the ACMG/AMP variant interpretation guidelines for genetic hearing loss.^10^ If only missense variants are calculated, the ratio is even lower (10%, 156/1493). The low ratio indicates that, with additional evidence, a significant number of the VUSs in this study, might await reclassification to P/LP. Nonetheless, the lower ratio can also be due to *GJB2* tolerance to genetic variation, consistent with the model adopted by the gnomAD database.^13; 33^, where the Z score of missense variants in the *GJB2* gene is −0.72, which signifies tolerance to variation.

The VUS burden observed in the *GJB2* gene raises significant challenges for clinicians and patients in clinical diagnostics. Although many studies related to *GJB2* have been published since it was first identified in 1997,^34^ the majority (72.8%) of the SNS variants on the coding region were still classified as VUS. The high rate of VUS in the *GJB2* gene is comparable with that of in the well-known tumor suppressor genes (*BRCA1/BRCA2*). Around 78% (8453/10854) of SNS variants in the *BRCA1/BRCA2* genes are recorded as VUS in the ClinVar (20210204 release). In order to sway a classification from VUS to P/LP or B/LB, further testing of family members or functional studies is required, which are time-consuming and labor-intensive and relies on family cooperation and consent.^35^ Another approach to minimize the number of VUS is data sharing. In this study, for example, we found at least two variants were classified as P/LP by the ClinGen HL-EP, which relies on an internal database. Nevertheless, long-term efforts are warranted to relieve the VUS burdens.

Medically significant discrepancies (upgrades to or downgrades from P/LP) in variant classifications between ClinVar and this study were observed in only 4% of variants with star 2+ and 12% for all. Most discrepancies were attributed to the lack or the availability of allelic data or functional data in the literature (Table S2). Considering that ClinVar is an open platform, subjective interpretations from different submitters inevitably generate inaccurate or conflicting classifications.^36^ Our variant interpretation and expert curation procedure provides accurate and reliable classifications, which reduced the number of conflicting interpretations. More importantly, we discovered that 54% (84/156) of P/LP missense variants were absent in ClinVar. Submitting these clinically actionable variants to ClinVar serves as valuable updates to the scientific and patient community and aids precise genetic counseling.

Analyzing the distribution of pathogenicity of missense variants across nine structural motifs helps us better understand the variant architectures of Cx26 and might assist us in determining critical regions where the PM1 could be applied. We found all the five residues (V43–E47) of the 3_10_ helix harbor P/LP missense variants, which could explain its important function. Residues of the 3_10_ helix are responsible for Ca^2+^ coordination^37^ and inter-protomer and intercellular interactions.^7^ More specifically, W44C and D46E inhibit the intercellular function of Cx26 wild-type, confirming the dominant-negative effect.^38; 39^ G45E displays increased hemichannel activity forms resulting in a gain-of-function effect.^40^ E47K and E47Q have defects in channel permeability resulting in the loss of gap junction function.^41^ Although recessive P/LP missense variants were enriched in TM2 domain, they were not equally distributed, thus challenging to the application of PM1 in this domain. Most P/LP variants were clustered in the TM2 region proximal to the E1 domain rather than the CL domain. This may be explained by the functional role of the two domains. E1 is critical to protein-protein interactions, whereas CL is a intracellular loop with disordered residues.^7^ To elaborate the critical region for PM1 activation in TM2 domain, more functional studies are warranted.

In addition, we found the cytoplasmic loops (CL and CT) are tolerant to variation (Table 1). Structurally, CL and CT are intracellular loops with disordered residues.^7^ Computationally, REVEL predicted only 5/100 and 4/62 missense variants on CL and CT are damaging.^42^ The data suggest that a variant located on CL and CT is most likely benign. Nevertheless, this work provides a new path for genetic professionals to define critical regions, although comprehensively interpreting a large number of missense variants in a gene is labor-intensive.

Variants in the *GJB2* gene cause considerable phenotypic heterogeneity. In this study, E1 and NT were found in favor of syndromic and dominant nonsyndromic phenotypes. Syndromic and dominant nonsyndromic phenotypes are usually caused by a gain of function mechanism where the integration of mutant Cx26 into homomeric or heteromeric connexon blocks their interactions with connexon in adjacent cells and finally produces structurally incomplete channels.^43; 44^ E1 has the most residues mediating inter-protomer interactions and inter-connexon interactions.^7; 43^ Moreover, six conserved cysteine residues that are critical for connexin docking and formation of functional gap junction channels are exclusively located on E1 and E2 (three in each)^45; 46^ Therefore, it is reasonable that E1 is enriched with syndromic and dominant nonsyndromic phenotypes. Regarding the favor of NT with KID syndrome, this could be explained that NT is crucial for the status (open or close) of gap junction channels.^7^ Mutations on the NT would increase the likelihood of aberrant hemichannel opening. KID syndrome is thought to result from channels that constantly leak ions, which impairs the health of the cells and increases cell death.^47^

The mode of inheritance should be carefully considered in variant interpretation, particularly for a disease with genetic and phenotypic heterogeneity. In this study, we noted two scenarios. The first is that a variant can be inherited in both autosomal recessive and autosomal dominant manner. To avoid erroneous classifications, we proposed that criteria supporting different inheritance patterns should not be activated simultaneously for a given classification. The second is the dual mode of inheritance on an amino acid residue. This scenario greatly impacts the activation of PM5 in variant interpretation. We refine the application of PM5 based on the inheritance model (Figure 4). In the *GJB2* gene, we found 13% of amino acid residues have a dual mode of inheritance, resulting in two different phenotypes. Although ClinGen HL-EP set different cutoffs of population data criteria (BA1, BS1, and PM2) for autosomal recessive and autosomal dominant hearing loss,^10^ further refinements are needed to accommodate the above, or similar, scenarios.

With the availability of classifications of all SNS variants in the coding region of the *GJB2* gene, we can estimate the genetic risk of DFNB1A amongst different populations. Although the estimated prevalence of DFNB1A displays considerable variability between populations, they are all comparable with the observed prevalence. Surprisingly, the high prevalence is contributed by a limited number of P/LP variants in most ethnicities. These findings are of importance for designing a limited, population-specific or pan-ethnics, cost-effective genetic screening panel for congenital hearing loss in newborns.^48^ That is, screening a limited number of variants could result in a low residual risk of DFNB1A and overcome the ethnic bias of a limited screening panel.^48^

In summary, we comprehensively interpreted all possible SNS variants in the coding region of the *GJB2* gene and characterized the distribution of pathogenic missense variants in this gene along with relevant genotype-phenotype correlations. Our results enhance the clinically adopted variant interpretation guidelines, which facilitate accurate genetic diagnoses of patients with *GJB2*-related deafness. The interpretation of *GJB2* SNS variants in the coding region also provides a prototype for genes with similarly high genetic and phenotypic heterogeneity.

## Supplemental Information description

Supplemental data include three figures and seven tables and can be found with this article online.

## Acknowledgments

No funding to declare.

## Declaration of Interests

Jiale Xiang, Xiangzhong Sun, Nana Song, Lisha Chen and Zhiyu Peng are employed at BGI Genomics at the time of submission. The other authors declare no conflict of interest.

## Web Resources

VIP-HL: https://hearing.genetics.bgi.com/

ClinVar: https://www.ncbi.nlm.nih.gov/clinvar/

gnomAD: https://gnomad.broadinstitute.org/

OMIM: https://www.omim.org/

HGMD: http://www.hgmd.org/

PubMed: https://pubmed.ncbi.nlm.nih.gov/

