## Supplemental Figures for "Comprehensive simulation and interpretation of single nucleotide substitutions in *GJB2* reveals the genetic and phenotypic landscape of *GJB2*-related hearing loss"

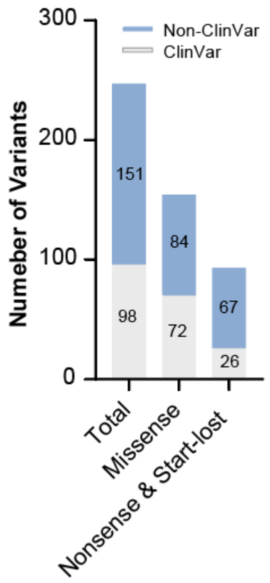


**Figure S1. ClinVar Inclusion of Pathogenic/likely pathogenic SNS variants.**
The number of different type of pathogenic/likely pathogenic SNS variants with or without record in ClinVar (20210204 release).

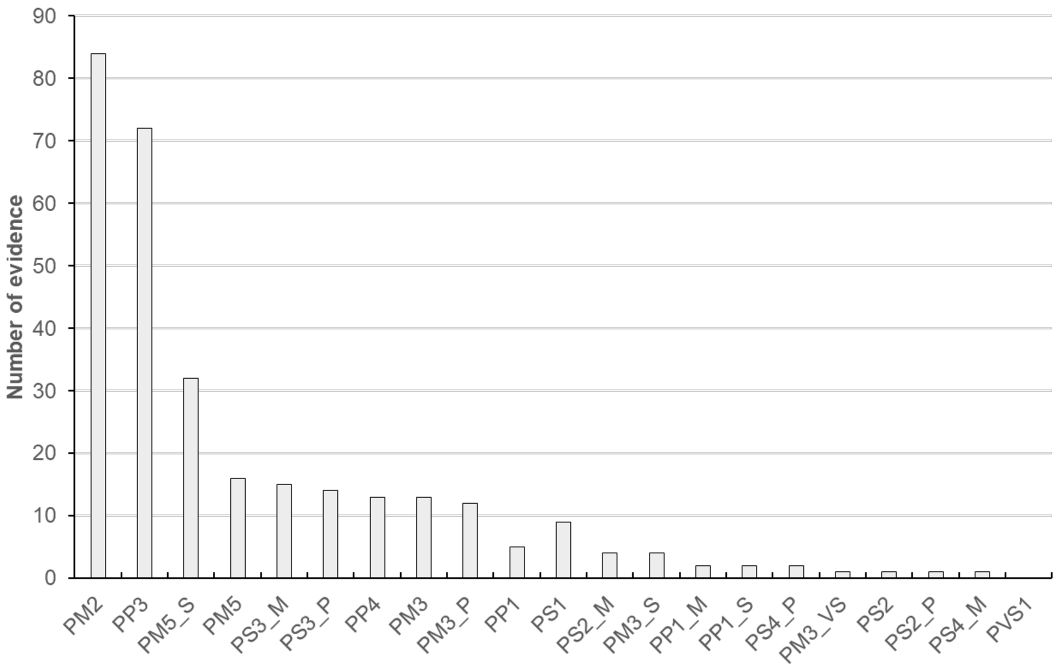


**Figure S2. Activated criteria for 84 non-ClinVar pathogenic/likely pathogenic missense variants.**

Evidence strength level: VS, very strong; S, Strong; M, moderate; P, Supporting.

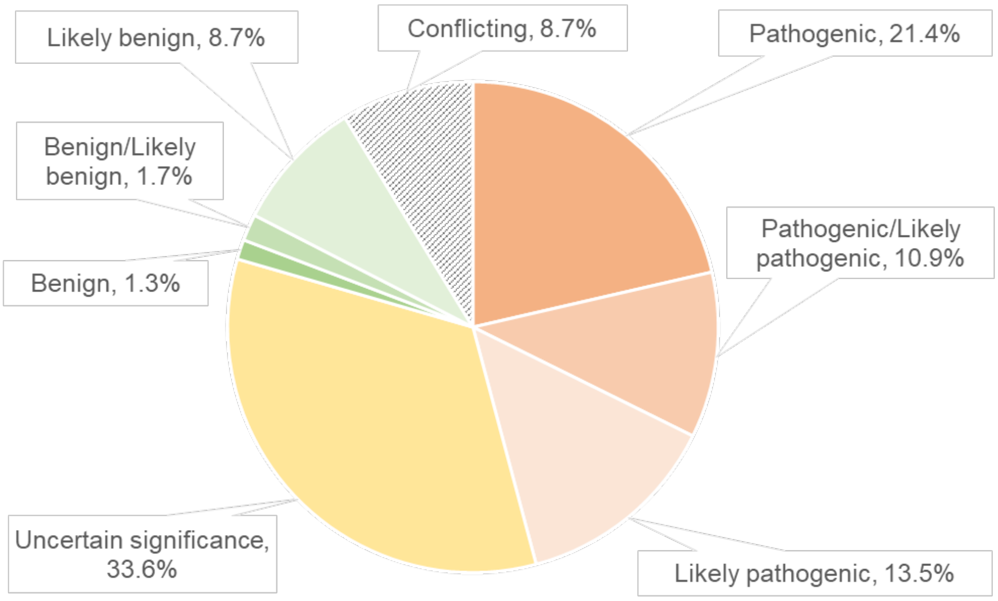


**Figure S3. *GJB2* SNS coding variants in ClinVar.**
Distribution of the classifications of 229 single-nucleotide substitution variants in the ClinVar database (20210204 release).
